# Soil communities following clearcut and salvage harvest have different early successional dynamics compared with post-wildfire patterns

**DOI:** 10.1101/2024.11.10.622867

**Authors:** Teresita M. Porter, Dave M. Morris, Emily Smenderovac, Erik J.S. Emilson, Lisa Venier

## Abstract

Understanding the impacts of harvest and subsequent silviculture practices at stand scales on the below-ground biota, and their associated nutrient cycling processes, is needed to more fully evaluate the sustainable management of boreal forest systems. While stand replacing wildfire is the primary natural disturbance mechanism in jack pine-dominated boreal forest systems; clearcut harvest also results in stand renewal so is sometimes used in silvicultural systems to emulate natural disturbance and renewal processes. In this study, we simultaneously assessed the successional trajectories of three major taxa of the below ground soil community, bacteria, fungi, and arthropods using DNA metabarcoding. The objectives of this study were to use a chronosequence framework to: 1) assess whether the soil communities following clearcut harvest and wildfire converge along a successional gradient, 2) assess when the soil community recovers following clearcut harvest to the pre-disturbance, mature, wildfire reference condition, and 3) assess the effects of cumulative disturbance on soil community succession (i.e., wildfire followed by salvage harvesting of fire-killed trees). We found that richness (alpha diversity) did not illustrate any clear patterns of convergence and could, therefore, underestimate recovery times, especially for soil arthropods. Comparisons of the underlying community composition (beta diversity) proved to be more informative. In this case, we found that different soil taxa following clearcut harvest recovered on different timelines compared with succession following stand-replacing wildfire. In general, bacteria appear to be the first to converge to post-wildfire conditions followed by arthropods, however, fungi did not converge within the time frame of the chronosequence. This suggests that more extended periods are required to achieve complete recovery of the soil fungal community to the pre-disturbance condition. The cumulative disturbance associated with salvage harvest appeared to have a greater (compounded) effect on soil communities when compared with wildfire or clearcut harvest. This work showcases the performance of a scalable method for monitoring a diverse arrange of soil biota using DNA metabarcoding. In future work, tracking fungal and arthropod soil communities may provide more insights into the longer-term effects of current forest management practices and provide guidance when comparing alternative approaches.

**Open Research Statement:** Sequences have been deposited to the NCBI SRA under the GRDI-Ecobiomics project accession PRJNA565010 for the BioSample accessions SAMN26926703 - SAMN26926795 used in this study. The MetaWorks v1.9.3 multi-marker metabarcode bioinformatic pipeline is available from https://github.com/terrimporter/MetaWorks. The ITS classifier based on the ITS UNITE+INSD full dataset v8.2 and trained to work with the RDP Classifier (https://github.com/terrimporter/UNITE_ITSClassifier). The COI classifier v4 is available from https://github.com/terrimporter/CO1Classifier. The code used to produce figures, including infile and metadata files, will be available on https://github.com/terrimporter/Chronosequence_HarvestType.

## Introduction

In Canada’s eastern boreal forest, jack pine (*Pinus banksiana* Lamb.) is a commonly occurring species that grows in pure to nearly pure stands on coarse-textured, glaciofluvial outwash deposits and is a staple species that provides both sawlog and pulpwood feedstock (Rudolph and Laidly 1990). It is an intolerant, pioneer species well adapted to stand-replacing wildfire. For example, serotinous cones release seed following high temperatures associated with wildfire resulting in high density pine-dominated stands. Silvicultural systems that emulate these natural disturbance processes are used as a model for sustainable forest management (O’Hara and Ramage 2013).

Forest management strategies that integrate stand-replacing disturbance like full-tree clearcut harvest may emulate certain aspects of a stand-replacing event such as natural wildfire (e.g., landscape pattern emulation). Clearcut harvest removes the overstory, but retains the existing downed woody debris and forest floor (McRae et al. 2001). Wildfire, on the other hand, is a pyrogenic process that kills trees but retains an overstory structure and consumes a portion of the forest floor as well as existing downed woody debris. Another forest operation currently used is post-wildfire salvage harvest that results in the partial consumption of the forest floor and downed woody debris by the stand-replacing wildfire followed by the removal of the fire-killed overstory trees.

The motivation for salvage harvesting following wildfire is to ameliorate economic losses by salvaging wood volumes that would have otherwise been lost through decomposition or blowdown following the wildfire event (Lindenmayer et al. 2004; Sessions et al. 2004). Salvage harvesting is controversial, however, because studies have shown that post-wildfire salvage may act more like a cumulative disturbance (Lindenmayer et al. 2004; Donato et al. 2006; Lindenmayer and Noss 2006; D’Amato et al. 2011; Lindenmayer, Burton, and Franklin 2012; Thorn et al. 2018). Specifically, salvage harvesting has been shown to: 1) reduce the ecosystem benefits provided by natural disturbance, 2) have negative impacts on many taxa, especially saproxylic species that require dead or dying wood for some part of their life cycle, 3) impair ecosystem recovery, and possibly alter ecosystem stability (i.e., not returning to the pre-disturbance condition), and 4) may result in additive effects associated with two disturbance events happening within 2-3 years (Lindenmayer et al. 2004; Thorn et al. 2018).

Understanding the below-ground impacts of above-ground harvest and silviculture relative to wildfire is needed to more fully evaluate the sustainable management of boreal forest systems (McRae et al. 2001). Examples of post-disturbance recovery dynamics that occur below ground include nutrient cycling processes such as decomposition, mycorrhizal symbioses, and soil respiration (Visser 1995; LeDuc and Rothstein 2007; Yermakov and Rothstein 2006; LeDuc et al. 2013). Reference information on soil biotic succession after natural wildfire disturbance can then be used to evaluate and compare the convergence and recovery trajectories after anthropogenic disturbances such as clearcut harvesting or salvage logging. Though there have been many studies that have examined the effects of fire and harvest disturbances on above-ground communities such as plants and animals (Schieck and Song 2006; Zwolak 2009; Kalies, Chambers, and Covington 2010; Clark and Covey 2012; Duguid and Ashton 2013; Fedrowitz et al. 2014; Gerstner et al. 2014; Michał and Rafał 2014; Alba et al. 2015; Willms et al. 2017; Vasconcelos, Maravalhas, and Cornelissen 2017; Lee 2018; Thorn et al. 2018; Carbone et al. 2019), less is known about below-ground communities largely because they tend to be difficult to study using conventional methods (Horton and Bruns 2001; Dooley and Treseder 2012; Holden and Treseder 2013; Dove and Hart 2017; Certini et al. 2021; Kudrin et al. 2023; Liu et al. 2023). Advances in molecular biodiversity monitoring, high throughput sequencing of signature DNA regions, also known as DNA metabarcoding, provides a rapid means for surveying the diverse soil organisms from bulk soil samples such as bacteria, fungi, and soil arthropods, that play important roles in nutrient cycling and soil remediation following disturbance (Claridge, Trappe, and Hansen 2009; Neary et al. 1999; Menta and Remelli 2020; Filialuna and Cripps 2021).

In our previous study of soil biotic succession following stand-replacing wildfires, we used DNA metabarcoding to show that each successional stage had unique biotic communities (Porter et al. 2023). Here, we extend our original study by comparing our post-wildfire soil successional reference with soil bacterial, fungal, and arthropod succession following full tree clearcut harvest (60 year chronosequence) and salvage harvest (20 year chronosequence). We used the chronosequence approach, where space is substituted for time, to evaluate soil community succession along the successional sequence (i.e., stand development stages). The objectives of this study were to: 1) assess whether soil communities become more similar over time by systematically analyzing community composition at each stand development stage, between the natural wildfire reference versus clearcut harvest or salvage harvest sites, 2) determine if/when the soil communities recover and return to the post-wildfire mature stand development stage, the natural disturbance endpoint in stand development, and 3) assess the effect of salvage logging (i.e., cumulative disturbance) on early soil community development. Further, we examined how various soil chemical properties correlate with these community changes.

## Methods

### Study Site Descriptions

Emulation silviculture is the use of harvest methods that emulate natural disturbances such as wildfire (McRae et al. 2001). In this paper, we compare the below-ground successional patterns in post-wildfire and post-harvest sites. A total of 31 sites were included in this study (Figure 1a) including 14 wildfire origin stands, 11 full-tree clearcut harvest sites, and 6 post-wildfire salvage harvest stands. All sites are in a cold, continental climate region, with an annual mean temperatures of 1.0 – 1.8 °C, Growing Degree-Days (GDD) from 1112 to 1263, and annual precipitation ranging from 68 – 82 cm yr^-1^ (growing season precipitation from 41 – 49 cm yr^-1^). Soils for all sites had comparable parent soil materials, well-drained, outwash sands, with well-developed Orthic Humo-Ferric Podzolic profiles. Stands were jack pine-dominated (*Pinus banksiana*) with many having minor amounts of black spruce (*Picea mariana*), trembling aspen (*Populus tremuloides*), and white birch (*Betula paperifera*). Wildfire and harvested sites generally had understories dominated by mosses, while salvage harvesting sites had litter and lichen dominated understories. Most stands are classified as Site Class I (Plonski 1981), with estimated site index values ranging between 18-22 m at breast height age 50.

**Figure 1.**
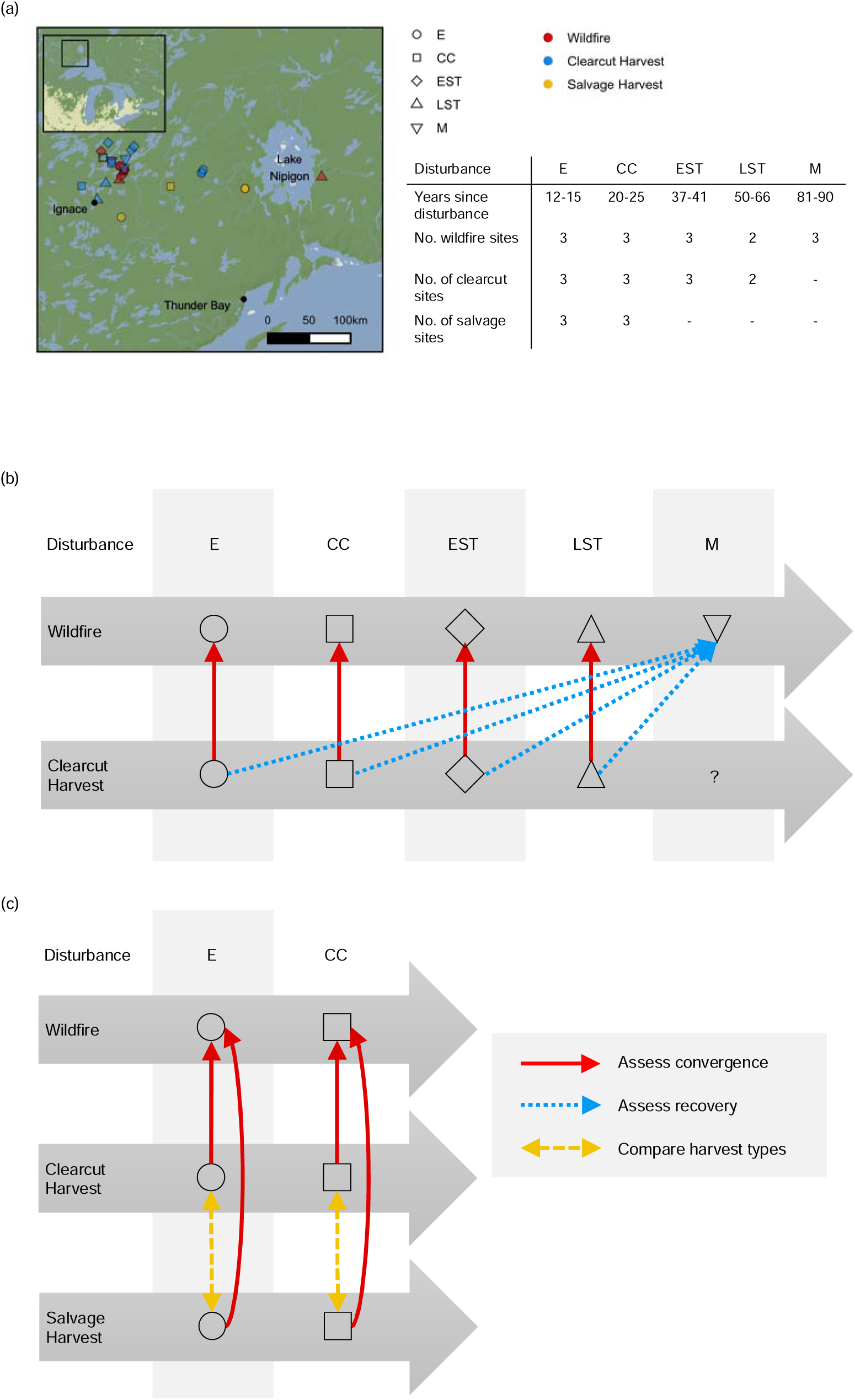
Sites and experimental design. (a) Inset map shows the Great Lakes Region, Ontario, Canada. Main map shows the distribution of sampling sites by disturbance type and stand development stage. Scale bar shows 50 km increments. Adjacent table summarizes the times since disturbance and the number of sites sampled for each disturbance type. (b) Summary of community composition comparisons at each stand development stage to assess whether sites become more similar over time (convergence) and comparisons with the reference wildfire condition at the mature stand development stage to assess whether sites return to the natural post-wildfire successional endpoint (recovery). (c) Summary of community composition comparisons with the wildfire reference condition and comparison of harvest types at each stand development stage. Abbreviations: Establishment (E), crown closure (CC), early self-thinning (EST), mature (M). © Stadia Maps © OpenMapTiles © OpenStreetMap © Stamen Design.

Each of the wildfire and salvage sites experienced spring, stand-replacing fires with Fire Weather Index (FWI) values ranging from 17.8 to 44.6 (exceeds extreme) in wildfire sites and 12.5 to 44.6 (exceeds extreme) in salvage sites. These sites were grouped along a time-since-disturbance sequence that correspond to stand development stages for jack pine that were comparable between the three disturbance types (establishment: 0-15 years, crown closure: 16-30, early self-thinning: 31-45, late self-thinning: 46-70, mature: 71+). Clearcut harvesting has only been operational for the past approximately 70 years, so our sites only have establishment to late self-thinning stages for this disturbance type. Post-wildfire salvage operations have only become commonplace in the last 20 years in Ontario’s northwest region, so our sites only have establishment to crown closure stages for this disturbance type.

A chronosequence is a space for time substitution approach that is often used when the time frame of the study question exceeds the life span of investigators. This approach is suitable for studying succession across decadal timescales when sites at different ages are following the same successional trajectory (Dyck and Cole 1994). The underlying assumption of this approach is that the primary differentiating factor between samples within a chronosequence is time since disturbance (Dyck and Cole 1994). To conform to the assumption noted above, our principal selection criterion was based on similarity in the basic site characteristics (*e.g.*, climate, soils, tree species composition, and productivity) described previously in more detail (Porter et al. 2023). Sites were jack pine-dominated (*Pinus banksiana*) separated by at least 1 km and represent independent stands, though some were the result of the same large-scale wildfire events.

### Soil Sampling and Laboratory Analysis

The field sampling design was previously described in detail (Porter et al. 2023). Briefly, in 2017, 3 replicate bulk soil samples were collected per field site. The bryophyte, organic LFH (litter, fermentation, and humus), and upper mineral soil layers were kept separate. Samples were immediately frozen at −20°C after sampling and held at that temperature until analysis. Additional samples of the LFH and mineral soil horizons (Ae, Bf collected separately) were also collected for pH and chemical determinations as previously described (Table S1).

### Molecular biology methods and sequencing

The molecular biology and sequencing methods were previously described in detail (Porter et al. 2023). Briefly, DNA extractions performed by the Natural Resources Canada lab at the Great Lakes Forestry Centre using the MoBio PowerSoil DNA Isolation Kit following the manufacturer’s instructions except that 200 ul of 100 mM AlNH_4_(SO_4_)_2_ was added to the tube along with soil and Solution C1 followed by a 10 minute incubation at 70°C to help lyse cells (Braid, Daniels, and Kitts 2003). The bacterial community was enriched using PCR by targeting the 16S v4-v5 region by prefixing the 515F-Y and 926R primers with Illumina adapters (Parada, Needham, and Fuhrman 2016); the fungal community was enriched by targeting the ITS2 region by prefixing the ITS9 and ITS4 primers with Illumina adapters (Menkis et al. 2012; White et al. 1990). Invertebrate communities were enriched using two sets of primers targeting portions of the COI DNA barcode region. The F230R_modN marker was targeted by prefixing the LCO1490 and the 230R_modN primers with Illumina adapters (Folmer et al. 1994; Gibson et al. 2015); and the BE marker was targeted by prefixing the B and E primers with Illumina adapters (Hajibabaei et al. 2012). For each sample a PCR cocktail of 5 µl template DNA (5 ng/µl in 10 mM Tris pH 8.5), 0.5 µl each of 10 µM forward and reverse primer, 25 µl HotStar Taq plus master mix kit 2x, and 19 µl of sterile water. PCR cycling conditions were as follows: 95°C for 5 minutes followed by 30 cycles (16S) or 35 cycles (ITS) or 40 cycles (COI) of 95°C for 30 seconds, 50°C for 30 seconds, and 72°C for 1 minute, followed by a final extension of 72°C for 5 minutes (10 minutes for COI) and hold at 4°C. PCR products were visualized then purified using magnetic bead solution (Agencourt AMPure XP, Beckman Coulter Life Science, Indianapolis, IN, USA) according to Illumina’s protocol (Illumina 2013). Sample indexes were added by amplifying 5 μl of the purified PCR product with 25 μl of KAPA HIFI HotStart Ready Mix, 5 μl of each Nextera XT Index Primer (Illumina Inc., San Diego, CA, USA) and 10 μl of UltraPure DNase/RNase-Free Distilled Water for a total volume of 50 μl. Thermal cycling conditions were as follows: 3 min at 98°C, 8 cycles of 30 sec at 98°C, 30 sec at 55°C, 30 sec at 72°C, and a final elongation step of 5min at 72 °C. Indexed amplicons were purified, quantified using a Qubit dsDNA BR Assay Kit (Life Technologies), and combined at equimolar concentration. Paired-end sequencing (2 × 250 bp) of the pools was carried out on an Illumina MiSeq at the National Research Council Canada, Saskatoon. 15% PhiX was added to help compensate for low sequence heterogeneity on the plate.

### Bioinformatic methods

Bioinformatic methods were previously described in detail (Porter et al. 2023). Briefly, demultiplexed, paired-end Illumina reads were analyzed on the General Purpose Science Cluster at Shared Services Canada. Metabarcodes were processed using the MetaWorks v1.9.3 multi-marker metabarcode pipeline available from https://github.com/terrimporter/MetaWorks using the standard sequence variant workflow (Porter and Hajibabaei 2022). Reads were paired using SEQPREP v1.3.2 using the default parameters except that the Phred score quality cutoff was set to 20 and the minimum overlap was set to 25 bp (St. John 2016). Primers were trimmed using CUTADAPT v3.2 using the default parameters except that the minimum sequence length was set to 150 bp, Phred quality score cutoffs at the ends was set to 20, and the maximum number of N’s was set to 3 (Martin 2011). Reads were dereplicated and denoised using VSEARCH v2.15.2, setting the minimum cluster size to retain to 3, so clusters with only 1 or 2 reads were removed as noise (Rognes et al. 2016). The 16S metabarcodes were taxonomically assigned using the RDP Classifier v2.13 using the built-in 16S reference set (Wang et al. 2007). The ITS metabarcodes were taxonomically assigned using the RDP Classifier with a custom-trained reference set based on the UNITE v8.2, QIIME release for Fungi v 04.02.2020, available from https://github.com/terrimporter/UNITE_ITSClassifier (Abarenkov et al. 2020). The COI metabarcodes were taxonomically assigned using the RDP classifier with a custom-trained COI classifier v4 reference set available from https://github.com/terrimporter/CO1Classifier (Porter and Hajibabaei 2018). In total we sequenced 49,721,154 paired-end reads for this study. After read pairing, primer trimming, and other quality control steps to remove poor quality and artefactual sequences, we retained 10,077,185 reads clustered into a total of 114,968 sequence variants (SVs) (Table S2).

### Data analysis

All data analyses was conducted in RStudio 2021.09.0 build351 using R v4.1.1 (RStudio Team 2016; R Core Team 2018). Diversity analyses were carried out with the vegan 2.6-2 package (Oksanen et al. 2018). For each sample, sequence reads were rarefied using the ‘rarecurve’ function to ensure that sequencing depth was sufficient across samples (Figure S1). In subsequent analyses, additional rare SVs that comprise less than 0.01% of reads were removed to account for the possibility of tag switching events (Elbrecht et al. 2017). Also in all subsequent analyses, for each sample, sequences from each soil layer were pooled together so that we could assess changes in composite diversity. After this filtering step we retained 1,671 SVs comprised of 6,642,061 reads. SV richness was assessed using the ‘specnumber’ function. Normality was assessed visually using the ‘ggqqplot’ function from the ggpubr 0.4.0 library and the base R Shapiro-Wilk test ‘shapiro.test’ function, and either Wilcox or t-tests were used to compare samples against the reference condition (Kassambara 2020). “Holm” adjusted p-values are shown for multiple comparisons. Binary Bray Curtis (Sorensen) dissimilarity matrices were created using the ‘vegdist’ function with binary=TRUE (Oksanen et al. 2018). Non-metric multidimensional scaling (NMDS) plots were calculated using the ‘metaMDS’ function using 3 dimensions after running ‘dimcheckNMDS’ from the goeveg 0.5.1 package and setting trymax=100 (Goral and Schellenberg 2018). Stress was assessed using the ‘stressplot’ function. Dispersion was assessed using the ‘betadisper’ function and the base R ‘anova’ function.

Environmental variables were fit to the ordination using the ‘envfit’ function with 999 permutations, and only variables with a p-value < 0.05 were plotted. We checked for significant interactions between disturbance type and stand development stage using permutational analysis of variance (PERMANOVA) using the ‘adonis2’ function with 999 permutations. When checking for convergence, at each stand development stage, we conducted pairwise comparisons of clearcut harvest disturbance with the wildfire reference condition using the ‘pairwise.adonis’ function with the pairwiseAdonis 0.4 wrapper for multi-level pairwise comparisons using adonis2 with 999 permutations and ‘Holm’ corrected p-values (Martinez Arbizu 2017). When checking for recovery, for each major group of soil taxa, we conducted pairwise comparisons of each stand development stage with respect to the mature wildfire reference condition. All plots were created using ggplot2 3.4.4 and the map was created using ggmap 4.0.0 (Wickham 2009; Kahle and Wickham 2013).

## Results

In general, the soil community was taxonomically diverse, especially for bacteria and fungi that tended to be dominated by a handful of high-level taxa (Figure S2). Across all sites, common taxa included bacterial Alphaproteobacteria, Acidobacteria, Chintinophagia, Gammaproteobacteria, and Planctomycetacia; fungal Agaricomycetes and Leotiomycetes; and arthropod Insecta and Arachnida.

### Establishing the reference condition: Succession following wildfire

Soil bacterial, fungal, and arthropod succession as well as community assembly patterns following natural wildfire disturbance were previously described in detail (Porter et al. 2023). Briefly, we found that bacterial communities were the first to recover (i.e., return to a comparable, community structure similar to the mature, wildfire reference condition) following wildfire by the crown closure stage (∼ 20 years). Bacterial communities were found to have a large core community with many shared taxa across each stand development stage. Moreover, bacterial community shifts were significantly correlated with changes in pH, total nitrogen, and total organic carbon. Fungi and arthropods, however, shared smaller core communities, i.e. had fewer taxa in common across stand development stages. In contrast with bacterial communities, fungal and arthropod communities had more taxon-replacement from one stand development stage to the next, i.e., each stand development stage supported more unique biodiversity compared with bacterial communities. Soil fungi and arthropod community shifts were significantly correlated with changes in total nitrogen and total organic carbon. Bacterial, fungal, and arthropod stand development stage bioindicator taxa were detected.

### Assessing convergence and recovery between wildfire and clearcut harvest stands

Convergence was compared between full tree clearcut harvest stands with the wildfire reference condition at the same stand development stage, from establishment to late self-thinning. Convergence of SV richness varied by taxonomic group (Figure 2 (a)). At the establishment stage, the bacterial and arthropod communities had higher richness in clearcut harvest compared with wildfire sites; and the fungal community had statistically significant higher richness in clearcut versus wildfire sites. For fungi, richness at each stand development stage fluctuated and appeared to converge by the late self-thinning stand development stage. For bacteria and arthropods, richness was similar between clearcut harvest sites and the wildfire reference condition at each stand development stage from establishment to late self-thinning.

**Figure 2.**
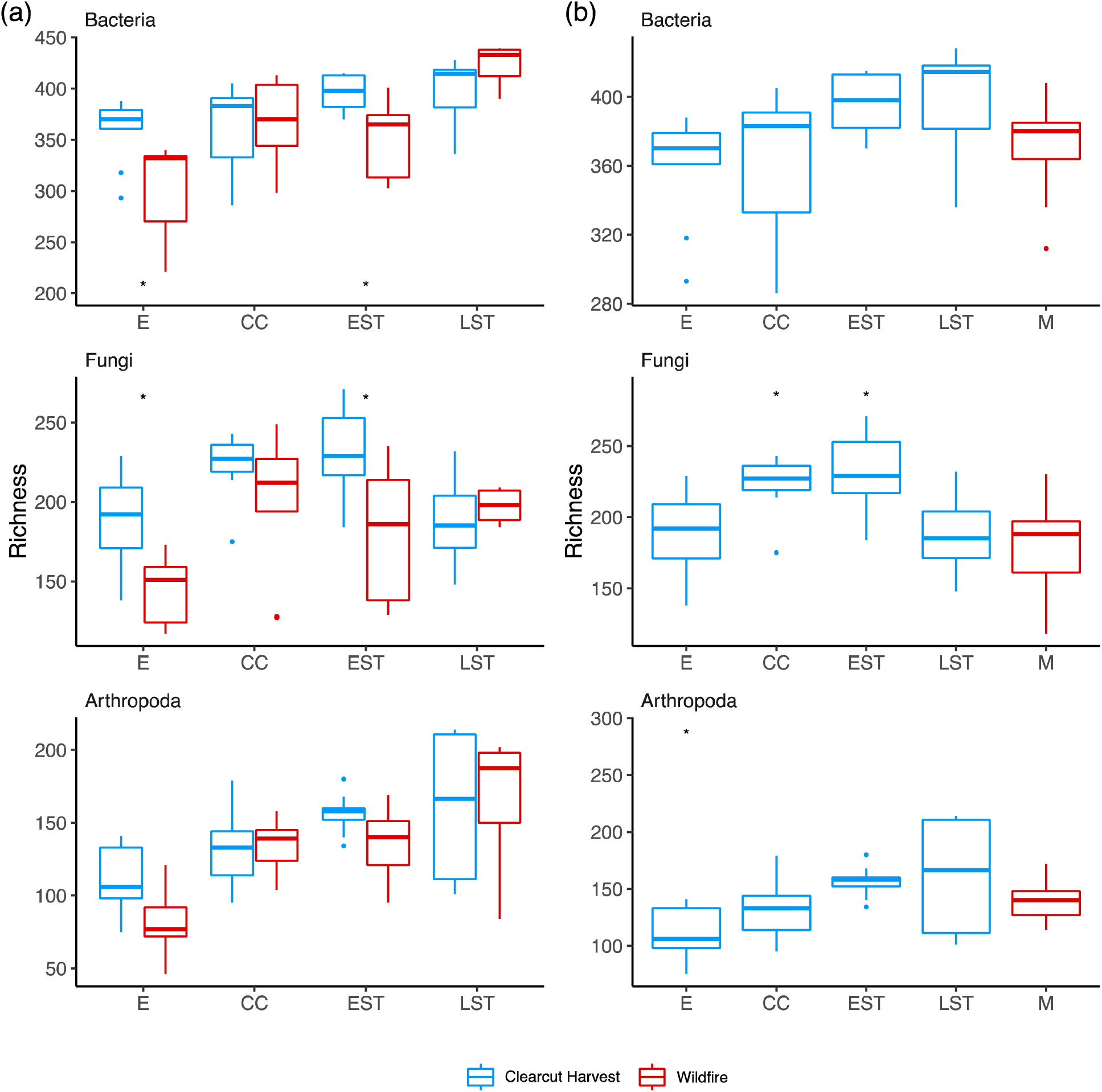
Analysis of richness did not reveal clear patterns of convergence and suggests resilience and early recovery times. Sequence variant richness was compared between the wildfire reference condition and full tree clearcut harvest sites to check for (a) convergence and (b) recovery across stand development stages. Wilcox (bacteria) or t-tests (fungi and arthropods) were used to compare soil communities and an asterisk is shown if the Holm corrected p-value was < 0.05. For part (a), statistical comparisons were grouped by taxon and stand development stage to compare richness between clearcut harvest and wildfire. For part (b), statistical comparisons were grouped by taxon to compare wildfire richness at each stand development stage with wildfire richness at the mature stand development stage. Abbreviations: Establishment (E), crown closure (CC), early self-thinning (EST), mature (M).

We also assessed recovery of clearcut harvest sites with respect to the mature, wildfire reference condition. Richness recovery varied by taxonomic group (Figure 2 (b)). For bacteria, richness in clearcut harvest sites was similar with the mature, wildfire reference condition across all stand development stages from establishment out to late self-thinning (adjusted p-values >= 0.05. For fungi, richness appeared to recover by the late self-thinning stand development stage. For arthropods, richness appeared to recover by the crown closure stand development stage.

Using beta diversity, instead of richness, allowed us to detect clearer patterns of convergence. Indirect gradient analysis was used to visualize convergence of beta diversity patterns following full tree clearcut harvest. NMDS plots show how community composition at each stand development stage shifted for each organismal group (bacteria stress = 0.08 and linear R^2^ = 0.98; fungi stress = 0.11 and linear R^2^ = 0.94; arthropod stress = 0.13 and linear R^2^= 0.92) (Figure 3 (a)). The asterisk indicates when communities are significantly different between the disturbance types, and when a drop in statistical significance occurs between successional stages (adjusted p-value > 0.05), it signifies when putative convergence in beta diversity occurred for each taxon. Based on this analysis, soil bacteria beta diversity appeared to converge at the crown closure stage, and this was driven by changes in pH and total organic carbon. For instance, at the establishment stage, wildfire bacterial communities were correlated with higher pH and lower total organic carbon, whereas clearcut harvest bacterial communities were correlated with lower pH and higher total organic carbon. By the crown closure stage, bacterial communities from wildfire and clearcut harvest sites appeared to be more similar and were similarly correlated with pH and total organic carbon. Soil fungal beta diversity from wildfire and clearcut harvest sites remained distinct across all stand development stages from establishment to late self-thinning (p-values < 0.05). For soil fungi at the establishment stage, wildfire communities were correlated with higher pH, whereas clearcut harvest communities were correlated with lower pH, higher total organic carbon and total nitrogen. Soil arthropod beta diversity appeared to converge at the early self-thinning stage. Similar to the other taxa, arthropod communities at the establishment stage following wildfire were correlated with higher pH, whereas clearcut harvest communities were correlated with higher total organic carbon and total nitrogen as well as lower pH. By the early self-thinning stage, soil arthropod communities in both wildfire and clearcut harvest stands appeared to be similar, with both being correlated with higher total organic carbon.

**Figure 3.**
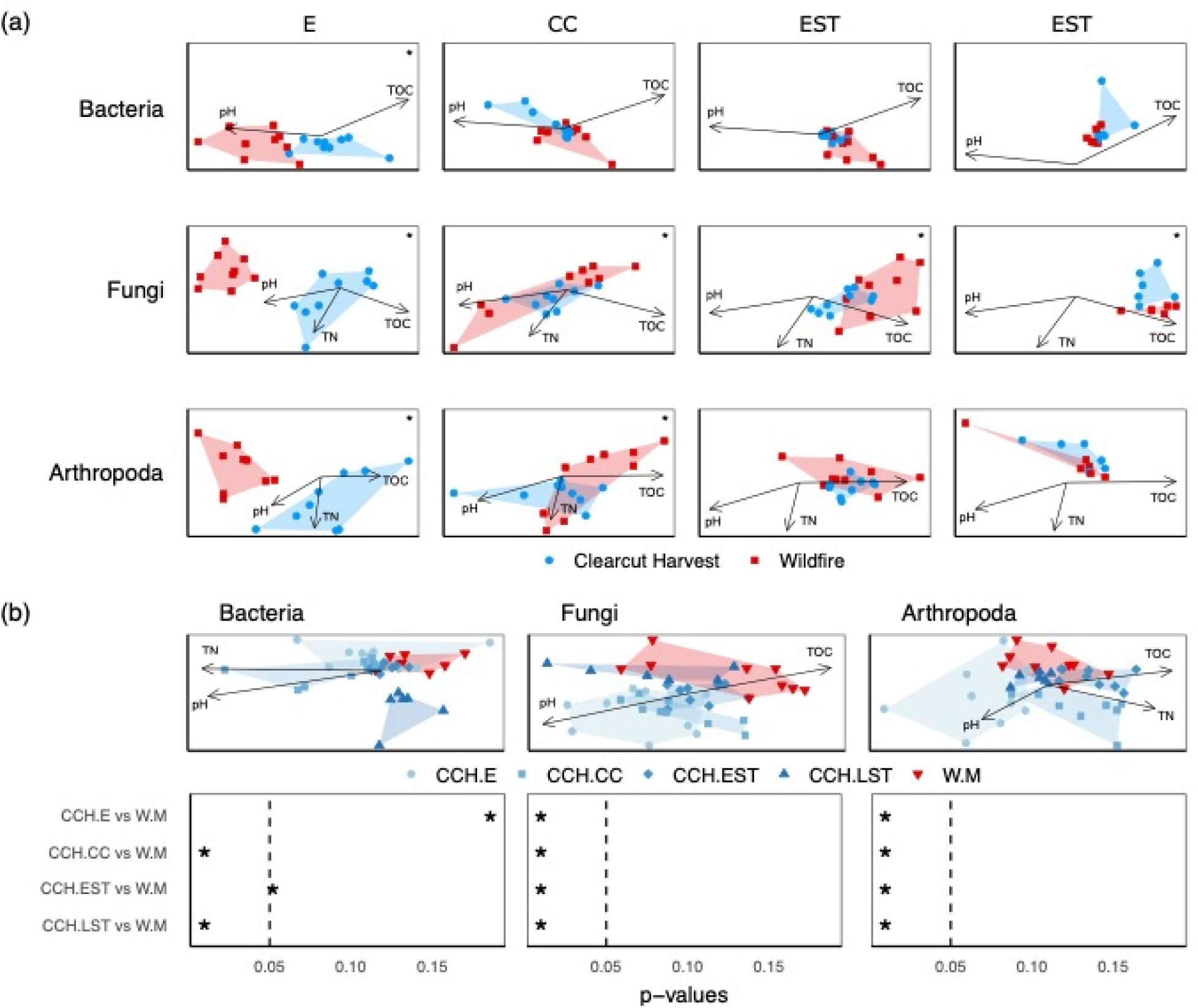
Analysis of beta diversity reveals clear patterns of convergence and shows a trajectory of recovery. Binary Bray Curtis dissimilarities (Sorensen distances) were compared between full-tree clearcut harvest sites and the wildfire reference condition. (a) Non-metric multi-dimensional scaling plots (NMDS) are used to assess community convergence at each stand development stage. Arrows indicate significantly correlated environmental variables with correlations > 0.30 and p-values < 0.05. Holm-corrected p-values < 0.05 from pairwise permutational analysis of variance (PERMANOVA) tests are marked with an asterisk. (b) NMDS plots are used to assess recovery trajectory, significantly correlated environmental variables, and the corresponding Holm-corrected p-values from pairwise PERMANOVA tests are shown below each plot. Abbreviations: Establishment (E), crown closure (CC), early self-thinning (EST), mature (M).

When beta diversity was used as a metric instead of richness, we detected later recovery or lack of recovery following clearcut harvest, depending on the taxa, when compared against the mature, wildfire reference condition. Indirect gradient analysis was used to visualize recovery of beta diversity. Recovery patterns are shown in Figure 3 (b) for each taxon (bacteria stress = 0.08 and linear R^2^ = 0.98; fungi stress = 0.12 and linear R^2^ = 0.93; arthropod stress = 0.14 and linear R^2^ = 0.89). Holm-corrected pairwise PERMANOVA p-values are shown below the recovery plots. Bacterial communities were distinct from the mature, wildfire reference condition at both the establishment and late self-thinning stages. Fungal and arthropod communities at each stand development stage appeared to be distinct from the mature, wildfire reference condition but the trajectory appeared to approach that of the mature, wildfire reference condition. All communities appeared to correlate with lower pH and higher total organic carbon over time, but arthropods were also correlated with lower total nitrogen over time. It is worth highlighting that because the variances in beta diversity were not homogenous, the significant differences found in the PERMANOVA could be due to this dispersion effect, especially for the fungal and arthropod communities as the clustering in the NMDS plots do suggest a recovery trajectory over time. For all taxa, variance in beta diversity was largely explained by stand development stage (28%, 22%, and 20%, respectively), followed by the interaction between stand development stage and disturbance type (full tree clearcut harvest and wildfire) (16%, 12%, and 11%) followed by disturbance type alone (4%, 5%, 4%) (p-value = 0.001).

### Comparison of full tree clearcut and salvage harvest versus natural wildfire disturbance

Soil diversity across two stand development stages, establishment and crown closure, were compared between wildfire, full tree clearcut harvest, and post-wildfire salvage harvest. In this case, the wildfire-origin sites were used as the reference condition at each stand development stage to assess the impact of harvest type. Using SV richness, all taxa showed convergence with the reference condition by the crown closure stage (Figure 4 (a)). Bacterial richness was not significantly different between harvest types (p-value > 0.05), but both were significantly elevated (p-value < 0.05) compared with the reference condition at the establishment stage. By the crown closure stage, bacterial richness was no longer significantly different from the reference condition (p-value > 0.05). Fungal richness was significantly elevated in clearcut and even more elevated in salvage harvest sites compared with the reference condition (p-value < 0.05) at the establishment stage. By the crown closure stage, fungal richness was no longer significantly different from the reference condition (p-value > 0.05). Arthropoda richness was not significantly different (p-value > 0.05) between harvest types but were significantly elevated (p-value < 0.05) compared with the reference condition at the establishment stage. By crown closure, arthropod richness was no longer significantly different from the reference condition (p-value > 0.05).

**Figure 4.**
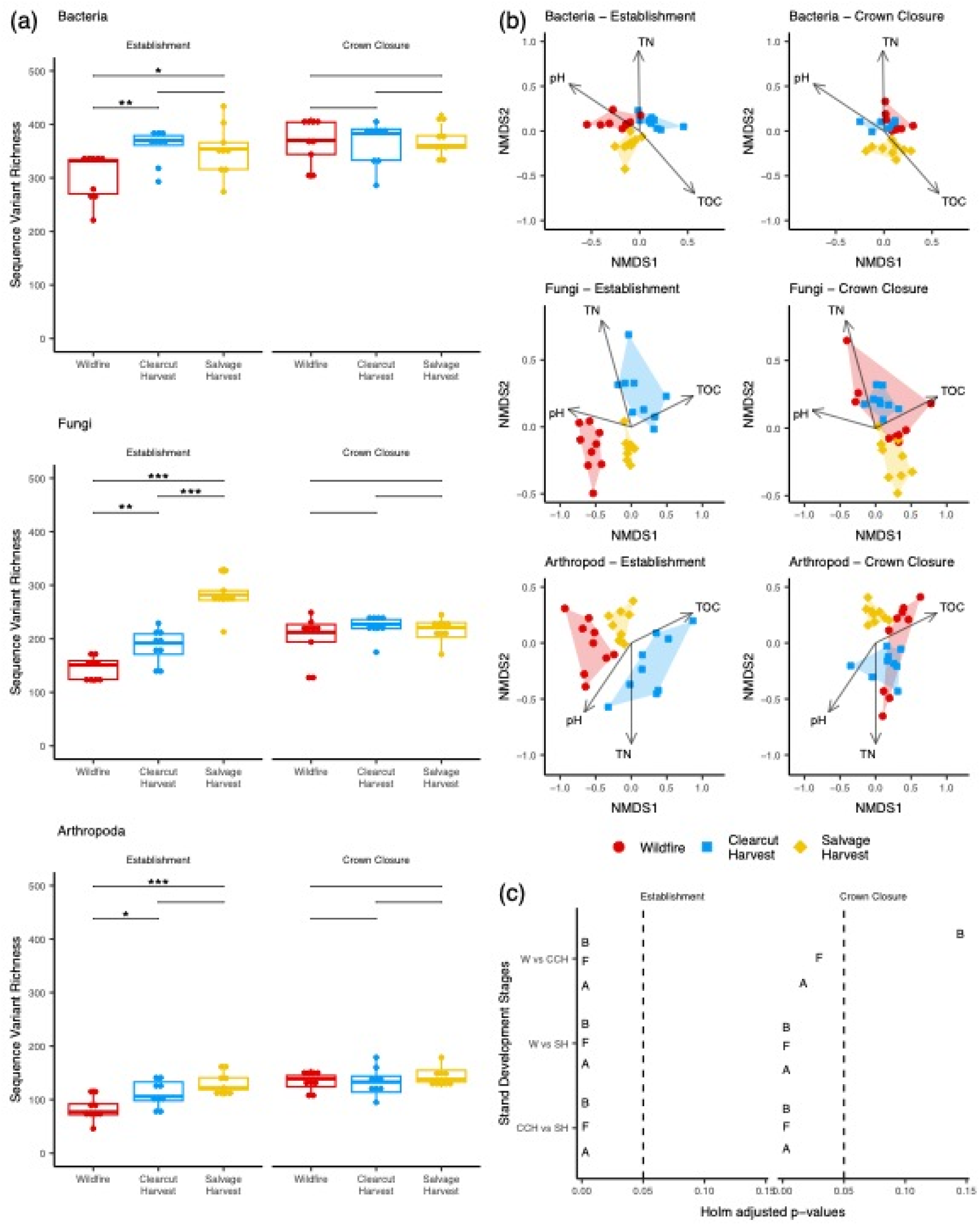
Salvage harvest appears to have a greater impact on soil community convergence than full tree clearcut harvest. The reference condition for clearcut and salvage harvest disturbances are wildfire samples in the establishment and crown closure stages. Convergence patterns are shown for (a) SV richness, (b) beta diversity comparisons among disturbance types using pair-wise PERMANOVA, and (c) indirect gradient analysis using NMDS with fitted environmental variables. Abbreviations: p-value >= 0.05 (NS), p-value < 0.05 (*), p-value < 0.01 (**), p-value < 0.001 (***); wildfire (F), harvest (H), salvage (S); bacteria (B), fungi (F), arthropoda (A); total organic carbon (g/kg) (TOC), total nitrogen (g/kg) (TN).

When examining beta diversity patterns, it was more difficult to detect convergence with the reference condition (bacteria stress = 0.07, R^2^ linear = 0.97; fungi stress = 0.11, R^2^ linear = 0.93; arthropoda stress = 0.13, R^2^ linear = 0.90) (Figure 4 (b)). At the establishment stage, higher pH in wildfire sites, and higher total organic carbon in clearcut harvest and salvage sites resulted in distinct bacterial, fungal, and arthropod groups. At both the establishment and crown closure stages, lower total nitrogen in salvage sties resulted in distinct bacterial, fungal, and arthropod groups. For all taxa, by the crown closure stage, wildfire and full tree clearcut harvest sites visually clustered more closely together compared to the salvage sites. Even after ∼ 20 years following disturbance, salvage sites clustered together for all taxa and appeared to form their own distinct group. For bacteria, these results were supported by PERMANOVA pair-wise tests shown in Figure 4 (c). For fungi and arthropods, PERMANOVA pair-wise tests showed that even at the crown closure stage, wildfire and fulltree clearcut harvest sites were significantly different but we feel this is largely due to significant dispersion in the variances across disturbance types for fungi (p-value = 0.004) and arthropods (p-value = 0.00000008). For all taxa, significant interactions were found between stand development stage and disturbance type with respect to variance in beta diversity (PERMANOVA, p = 0.001).

## Discussion

### Convergence and recovery of wildfire- and clearcut harvest-origin stands following stand-replacing disturbance

We assessed whether the soil biotic communities in stands disturbed by full-tree clearcut harvesting converged with wildfire disturbed stands and showed signs of recovery (i.e., tracking towards the mature, wildfire reference condition) on similar trajectories as naturally disturbed stands following wildfire. Assessment of convergence involved the comparison of post-wildfire and harvest origin stands at comparable stand development stages. We found that richness was numerically higher for bacteria and significantly higher for fungi in full tree clearcut harvest sites compared with wildfire sites during the initial stand establishment stage (0-10 years following disturbance. This result corresponds with a previous study that assessed microbial diversity in post-wildfire and post-harvest jack-pine stands and found lower microbial diversity following wildfire (Staddon, Duchesne, and Trevors 1998). These initially higher richness values may, in part, be due to a legacy effect of retaining a portion of the pre-harvest bacterial and fungal communities suggesting some level of resilience.

Bacteria also showed beta diversity convergence with the post-fire community composition by the crown closure stage. The relatively early/rapid convergence of the bacterial communities suggests that bacteria are not overly sensitive to the demonstrated differences between clearcut harvesting and wildfire. This lack of sensitivity contrasts strongly with the other taxa. Arthropod communities did not converge until the late self-thinning stage and the fungal communities that did not show convergence. These results indicate that harvesting disturbances to soil fungal and arthropod communities are not directly analogous to those experienced in typical fire successional dynamics and that clearcut harvesting of mature forests does create different habitat conditions across most of the stand development stages. These differences in response between taxa are likely driven, in part, by the provision of different habitat conditions created by these two disturbance types that could represent distinct successional processes between these two disturbances.

As soil biotic communities have been demonstrated to be highly responsive to changes in physicochemical conditions (e.g., moisture, temperature) (Wilhelm et al. 2017), these soil biotic community differences are likely reflecting real differences in site conditions. For example, wildfire consumes a substantial portion of the pre-disturbance forest floor and downed woody debris (DWD) complex, volatilizes N, raises pH via ash deposition, and the blackened soil surface absorbs more incoming solar radiation. In contrast, forest floors and existing DWD following harvesting largely remain intact, although some mechanical disruption does occur along wood extraction trails. As highlighted in Figure 3, community differences between wildfire and harvested stands at stand establishment were driven by higher pH, but lower concentrations of TOC and TN on the wildfire sites.

We also assessed the recovery of soil biotic communities back to their pre-harvest, mature, fire-origin condition (i.e., natural succession endpoint). This assessment involved the comparison of harvested stands at each stand development stage against the mature, wildfire reference condition. The rapid recovery of bacterial communities in full tree clearcut harvested sites further supports the likelihood of a legacy effect where more organisms from the ‘mature’ (pre-disturbance) condition were retained through harvesting when compared against their wildfire disturbance counterparts. This legacy effect is also likely a function of retaining the existing forest floor and DWD material.

Our observed effects could also have implications for carbon sequestration. In Canadian old boreal forests, it was shown that soil carbon pools began to decrease, or at least showed no notable increases, in mature forests (Gao et al. 2018). They also showed that early successional stages accumulate the most tree carbon, many of the organisms that are plant pathogens or saprotrophs are present in mature soils, and so they could potentially limit carbon accumulation if they are more abundant in early successional clearcuts. In Canadian boreal forests in northern Ontario, despite measured declines in soil C pools following clearcut harvesting in jack pine and black spruce-dominated stands, it was found that these pools had returned to pre-disturbance levels within 20 years after disturbance (Morris et al. 2019).

### Impact of salvage logging relative to wildfire and clearcut harvest disturbances

In this study, soil biotic community composition across all three taxa in salvage-logged sites appeared distinct from either the wildfire or clearcut harvest sites and remain so through the crown closure stage of stand development (out to 20 years following disturbance). As highlighted in Figure 4 (b), these differences are largely driven by the low soil N concentrations on salvage logged sites. These differences may have implications for soil function including nutrient cycling in these coarse-textured, nutrient poor soils. A previous study that assessed the impacts of wildfire and salvage harvest suggested that soil nutrients in jack pine and black spruce stands would not likely return to their pre-burn level even in a 110-year rotation (Brais, David, and Ouimet 2000). In a review comparing wildfire and harvest systems, differences in nutrient pools between wildfire and other stand-replacing disturbances such as clearcut harvest, or salvage harvest following wildfire were also highlighted (McRae et al. 2001). In some cases, it has been suggested that harvesting of trees from sites before they have recovered from a natural disturbance, may represent a second disturbance resulting in compounded effects that, in turn, result in longer-term alteration of the affected biotic communities (D’Amato et al. 2011; Paine, Tegner, and Johnson 1998; Maynard et al. 2014) which is consistent with what we observed here. Post-fire salvage logging has been shown to reduce the habitat preferred by pyrophilous species such as certain arthropods and fungi (Lindenmayer and Noss 2006; Cobb et al. 2011; Pérez Izquierdo et al. 2021). In contrast, responses of fungi may be less severe than reductions in arthropods, as ectomycorrhizal fungi can persist on the roots of surviving trees after wildfire or as resistant structures in the soil, after a severe fire or salvage logging where most of the trees are killed or removed (Visser 1995). Reductions in fungi, however, can have ecological consequences that are perceived as positive or negative for forest operations. Some fungi are important recyclers of organic material, or plant mutualists, but others are adapted to cause plant disease. Where this legacy fungal inoculum is reduced, by salvage logging for example, development of soil communities may be slow, or altered, as recolonization proceeds primarily from spores dispersed by air, animal, or soil (Dahlberg 2002; Baar et al. 1999; Glassman et al. 2016). Slow spore dispersal rates might explain the lack of fungal convergence between wildfire and salvage sites out to 20 years post-disturbance. Similarly, the removal of fire-killed trees during salvage harvest, reduces the downed woody debris pool necessary for some elements of the arthropod community such as beetles and flat bugs, contributing to the observed community differences (Cobb et al. 2011; Cobb, Langor, and Spence 2007; Heikkala, Martikainen, and Kouki 2017). The removal of these tree residues can also impact soil conditions by changing site microtopographic characteristics like temperature and moisture that can impact soil microbial communities (Pereira et al. 2018). It has been proposed that the retention of standing trees and deadwood can moderate observed changes in the fungal community (Mayer et al. 2022; Prescott 2024). There is also evidence that compounding disturbances in forest management practices can contribute to impacts to the arthropod community (Cobb, Langor, and Spence 2007; Work et al. 2023). Our results suggest that salvage logging can extend the time for convergence of the forest soil biotic community towards the wildfire disturbance condition and creates unique communities that are not part of the natural successional sequence. Although our timelines were limited here, salvage logging appears to alter the successional pathways of the soil communities which undermines the assumptions of the natural disturbance emulation approach.

### Management Implications

In many jurisdictions, including Ontario, Canada, forest sustainability includes reference to using forest practices that emulate natural disturbances (END) to conserve biodiversity and their associated ecological processes (i.e., Ontario’s Crown Forest Sustainability Act of 1994). In the boreal forest, END has generally focused on landscape pattern emulation (e.g., the size and distribution of harvest relative to wildfire). However, at the stand/site scale, wildfire and harvest are not the same disturbance mechanism. One is a pyrochemical process that kills, but retains, the overstory tree stems, but consumes a substantial portion of the existing forest floor and downed woody debris. In contrast, clearcut harvesting is a mechanical process that removes the merchantable overstory trees but leaves the forest floor and DWD complex largely intact. Even though both are stand-replacing disturbances that reset the disturbed areas to the establishment stage of stand development, the conditions immediately after disturbance are different. Within the context of END, the underlying assumption is that there will be convergence over a reasonable period of time and will eventually resemble (recover) the pre-disturbance condition, that, in turn, would define the “ecological” rotation (Kimmins 1974; Prescott 2024).

The lack of convergence of arthropod and fungi communities seen in this study does raise a concern with the END paradigm. Our results suggest that soil communities in harvested sites can remain significantly different through successional stages up to the late self-thinning stage (46-70 years following disturbance). Given the spatial extent of clearcut harvesting, these results suggest potential large-scale changes to the distribution of distinct soil communities across Ontario’s boreal forest. The use of prescribed fire, as a site preparation tool, may be one approach that could be used to ameliorate these effects, but in general they are difficult to implement and not commonly done. Introducing some of the aspects of fire disturbance with additions such as wood ash from bioenergy production may be another option to reducing the gap between harvest and fire origin soil communities (Hannam et al. 2019).

The lack of recovery in both the fungi and arthropod communities out to the late self thinning stage would also suggest that timber extraction rotations, for at least a portion of our managed stands, need to be longer than 70 years, and not based primarily on the culmination of MAI (mean annual increment), to allow for the full recovery of the soil biotic community. This is consistent with previous forest management recommendations and would ensure representation of old forests conditions through both retention and succession (Prescott 2024). Unfortunately, we did not have any harvest origin stands beyond 66 years to confirm when full recovery is achieved. We do know from our previous study of post-wildfire soil biota succession that soil communities in fire origin stands remained unique out to at least age 80 (Porter et al. 2023).

Results for both convergence and recovery highlight the value of using beta diversity rather than just species richness (alpha diversity) to assess successional trajectories as it provides much more information relevant to decision making for forest management. In addition, the beta diversity results demonstrate clear distinctions between the response of individual taxa highlighting the value of a multi-taxonomic approach to sustainability assessments (Porter et al. 2023; Venier et al. 2014). Beyond beta diversity, however, we also need to understand how these shifts in community composition affect soil function.

### Methodological Considerations and Future Direction

Adaptive management is an approach that recognizes that policy should be treated as a testable hypothesis and its effectiveness should be measured to support policy refinement.

Biodiversity is an essential indicator of forest management policy to achieve sustainability. Our work highlights the capacity for DNA metabarcoding to inform forestry-related questions about below-ground biodiversity that are generally challenging to study using more conventional methods. The metabarcoding approach also supports the inclusion of a wide diversity of taxa in the assessment. Our results clearly demonstrate the need for multi-taxonomic assessments due to the taxonomic variation in response to disturbances. The results also highlight that a wider array of taxa provides more insight at functional levels that should be examined in the future. Because a major limitation when working with such a broad array of organisms is lack of representation in reference sequence databases at a fine taxonomic level, this work focused on the analysis of sequence variants. Future studies using similar techniques could be used to further assess the potential ecological implications of the differences observed in this study. One potential focus could be investigating the aboveground-belowground linkages facilitated by organism exchange and, potential functional activities identified with enzyme assays or functional databases. This is an important avenue for future research and could involve the use of functional databases such as FUNGuild, Faprotax, BETSI, GLoBi that can map trophic interactions, enzyme assays that measure changes in potential nutrient use, or shotgun sequencing that can provide information on the functional genes that are present (Tringe et al. 2005; Jackson, Tyler, and Millar 2013; Poelen, Simons, and Mungall 2014; Louca, Parfrey, and Doebeli 2016; Nguyen et al. 2016; Joimel et al. 2021). Additional analysis of these data can be used to assess the functional dynamics of recovery and understand which components of the community are responsive or resilient to the changes represented by the different forest age classes, and even which organisms can colonize from external sources, versus those that require *in-situ* regeneration.

Tests of the effectiveness of the natural disturbance emulation policy are rare and difficult. This study offers a unique comparison of biodiversity response to harvest relative to fire. The chronosequence approach provides unique information about the contrast between harvest and fire over time frames that are not achievable in long-term studies. Efforts to establish and maintain chronosequence studies should be a focus of effectiveness research, especially as the information necessary to establish chrono-sequence studies is being lost over time. Also, in the Ontario boreal forest, we are just now seeing post industrially-harvested stands reaching 70 years and so there is significant potential to extend these chronosequences over longer timeframes to provide additional insights with respect to full recovery and ecological rotations.

## Conclusions

This chronosequence study combined with DNA metabarcoding of soil biodiversity has provided some unique insights into the effectiveness of natural disturbance emulation as a forest management paradigm. Soil fungi and arthropods were the most responsive taxa indicating that there are significant differences in soil community composition through all examined stand development stages, and that some taxa do not recover to the mature, fire-origin reference condition over a 60-year timeframe. A comprehensive look at potential functional differences is now needed to better understand the ecological implications of these community differences. Our results do, however, suggest that innovative management approaches should be considered to support soil communities that, in turn, support resilient forests. For example, we recommend consideration of use of prescribed burns as a site preparation tool, where feasible, or the addition of wood ash from bioenergy production to improve the emulation of wildfire. We also recommend allowing for longer rotations and balancing age class distributions for the provision of habitat for the full range of soil biota. Lastly, we recommend only limited use of salvage logging after fire because it shifts soil communities away from a natural fire-origin composition. Where conducted, salvage-logged areas should have extended rotations to account for the lag in convergence and recovery.

## Author contributions

DM and LV conceived of the project and conducted site selection. DM, LV and EE obtained funding for the project. LV and EE provided the laboratory services for sample processing. TMP did the bioinformatics, data analysis, and wrote the manuscript. ES, DM, and LV assisted with data analysis. All authors contributed to editing the manuscript.

## Acknowledgements

TMP was funded by the Government of Canada through the Genomics Research and Development Initiative (GRDI) Metagenomics-Based Ecosystem Biomonitoring (Ecobiomics) project. We are grateful to Derek Chartrand, Kerrie Wainio-Keizer and Susan Bowman at the Great Lakes Forestry Centre for soil sample processing and conducting DNA extractions; Marie-Josée Morency and the Seguin lab for amplicon and sequencing library preparation; as well as to the Natural Research Council Canada sequencing facility in Saskatoon for Illumina MiSeq sequencing services.

