## SupplementaryMaterial for "Soil communities following clearcut and salvage harvest have different early successional dynamics compared with post-wildfire patterns"

### Supplementary Results

**Figure S1. Rarefaction curves plateau indicating sequencing depth was sufficient for each amplicon.** Across all samples in this study, the number of sequence variants (SVs) detected from bacteria (16S) is highest, followed by fungi (ITS), and least for arthropods (COI).

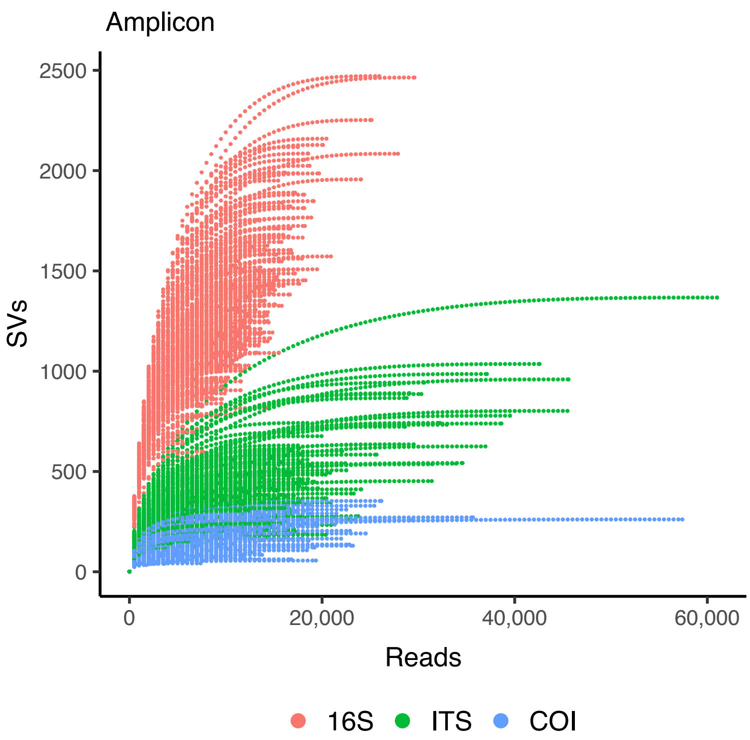

**Figure S2. Overview of taxa from wildfire, full tree clearcut, and salvage harvest sites.** Sequence variant richness (total number of unique SVs) is shown for (a) bacteria, (b) fungi, and (c) arthropods showing dominant taxa for each disturbance type at each stand development stage. The same dominant classes were also detected with relative read abundance (data not shown). Abbreviations: establishment (E), crown closure (CC), early self-thinning (EST), late self-thinning (LST), mature (M).

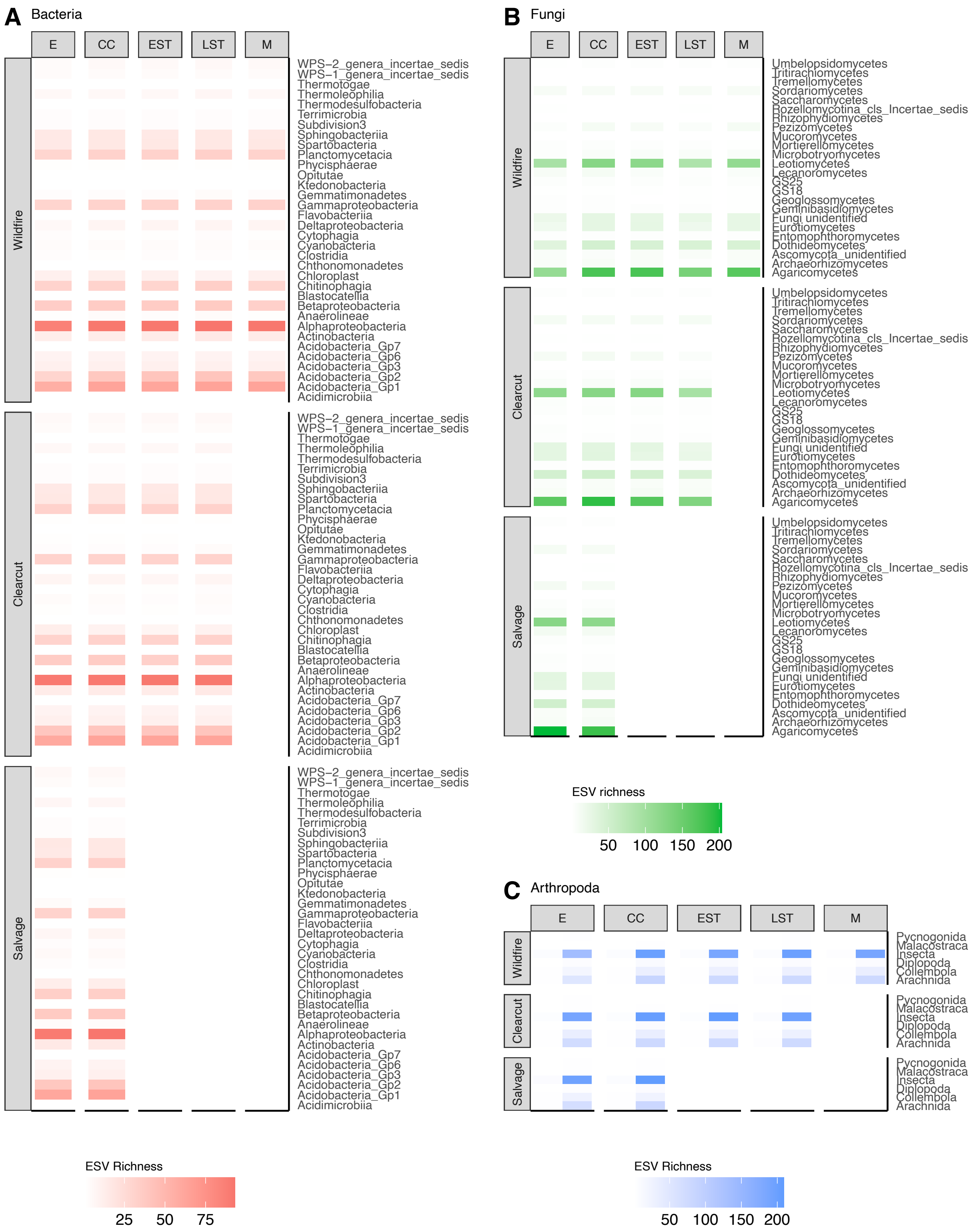
A Bacteria
E
CC
EST
LST
M
Wildfire
Clearcut
Salvage
WPS−2_genera_incertae_sedis
WPS−1_genera_incertae_sedis
Thermotogae
Thermoleophilia
Thermodesulfobacteria
Terrimicrobia
Subdivision3
Sphingobacteriia
Spartobacteria
Planctomycetacia
Phycisphaerae
Opitutae
Ktedonobacteria
Gemmatimonadetes
Gammaproteobacteria
Flavobacteriia
Deltaproteobacteria
Cytophagia
Cyanobacteria
Clostridia
Chthonomonadetes
Chloroplast
Chitinophagia
Blastocatellia
Betaproteobacteria
Anaerolineae
Alphaproteobacteria
Actinobacteria
Acidobacteria_Gp7
Acidobacteria_Gp6
Acidobacteria_Gp3
Acidobacteria_Gp2
Acidobacteria_Gp1
Acidimicrobiia
ESV Richness
25
50
75
B Fungi
E
CC
EST
LST
M
Wildfire
Clearcut
Salvage
Umbelopsidomycetes
Tritirachiomycetes
Tremellomycetes
Sordariomycetes
Saccharomycetes
Rozellomycotina_cls_Incertae_sedis
Rhizophydiomycetes
Pezizomycetes
Mucoromycetes
Mortierellomycetes
Microbotryomycetes
Leotiomycetes
Lecanoromycetes
GS25
GS18
Geoglossomycetes
Geminibasidiomycetes
Fungi unidentified
Eurotiomycetes
Entomophthoromycetes
Dothideomycetes
Ascomycota_unidentified
Archaeorhizomycetes
Agaricomycetes
ESV richness
50
100
150
200
C Arthropoda
E
CC
EST
LST
M
Wildfire
Clearcut
Salvage
Pycnogonida
Malacostraca
Insecta
Diplopoda
Collembola
Arachnida
ESV Richness
50
100
150
200

**Table S1. Overview of site characteristics for each disturbance type and stand development stage.**

| Disturbance type | Site characteristic | Stand Establishment | Crown Closure | Early Self-Thinning | Late Self-Thinning | Mature |
|---|---|---|---|---|---|---|
| Wildfire | Dominant understory | Moss, deadwood (fire origin), blueberry | Moss, deadwood (fire origin), bunchberry | Moss, litter | Moss, deadwood (self-thinning origin), mayflower | Moss, lichen, honey suckle |
| | Mean pH[a] | 5.76 ± 0.23 | 5.19 ± 0.48 | 5.04 ± 0.46 | 5.06 ± 0.38 | 4.97 ± 0.36 |
| | Mean total organic carbon (g/kg)[a] | 244 ± 180 | 333 ± 231 | 330 ± 231 | 336 ± 234 | 323 ± 225 |
| | Mean total nitrogen (g/kg)[a] | 8.58 ± 5.79 | 9.07 ± 7.30 | 7.65 ± 5.54 | 8.67 ± 6.01 | 7.81 ± 5.50 |
| Full tree clearcut harvest | Dominant understory | Blueberry, Honeysuckle, Moss, Bunchberry, Raspberry | Moss, Litter, Labrador Tea, Bunchberry, Blue-bead Lily, Blueberry | Moss, Litter, Bunchberry | Moss, Bunchberry, deadwood (self-thinning) | - |
| | Mean pH[a] | 5.24 ± 0.41 | 5.43 ± 0.26 | 4.82 ± 0.68 | 5.26 ± 0.24 | - |
| | Mean total organic carbon (g/kg)[a] | 306 ± 213 | 330 ± 229 | 334 ± 231 | 326 ± 230 | - |
| | Mean total nitrogen (g/kg)[a] | 8.35 ± 5.83 | 9.67 ± 6.78 | 9.89 ± 6.94 | 8.52 ± 6.14 | - |
| Salvage harvest | Dominant understory | Litter, Blueberry | Lichen | - | - | - |
| | Mean pH[a] | 5.56 ± 0.22 | 5.01 ± 0.37 | - | - | - |
| | Mean total organic carbon (g/kg)[a] | NA | 301 ± 214 | - | - | - |
| | Mean total nitrogen (g/kg)[a] | 7.22 ± 4.79 | 7.34 ± 5.20 | - | - | - |

Note: Extractable phosphorus was also measured but not included here because it was not significantly correlated with soil community shifts, neither in this study nor in our previous study focusing on succession in post-wildfire soils (Porter et al. 2023).

[a] Mean value pooled across soil layers with standard deviation.

Abbreviations: Not sampled (-), Not available (NA).

**Table S2: Reads and sequence variants (SVs) at each major bioinformatic step.**

| Step | COI-BE | COI-F230 | ITS2 | 16Sv4v5 | Total |
|---|---|---|---|---|---|
| **Raw reads** | 7,726,307 x 2 | 18,578,038 x 2 | 11,376,813 x 2 | 12,039,996 x 2 | 49,721,154 x 2 |
| **Paired reads** | 6,971,930 | 17,041,560 | 9,780,624 | 10,411,151 | 44,205,265 |
| **Primer-trimmed reads** | 6,014,076 | 16,650,830 | 9,567,562 | 10,208,799 | 42,441,267 |
| **Denoised SVs[a] (Reads)** | 1,260 (97,266) | 7,733 (2,741,702) | 27,884 (3,952,018) | 78,091 (3,286,199) | 114,968 (10,077,185) |

[a] Removal of sequence clusters with less than 3 reads, sequences with errors, putative chimeras, and obvious pseudogenes to create a set of sequence variants (SVs).
